# Insights into the ribosome function from the structures of non-arrested ribosome nascent chain complexes

**DOI:** 10.1101/2022.02.21.480960

**Authors:** Egor A. Syroegin, Elena V. Aleksandrova, Yury S. Polikanov

## Abstract

During protein synthesis, the growing polypeptide chain threads through the nascent peptide exit tunnel that spans the body of the large ribosomal subunit while simultaneously acting as a modulator of ribosomal activity by itself or by sensing various small molecules, such as metabolites or antibiotics appearing in the tunnel. While arrested ribosome nascent chain complexes (RNCCs) have been extensively studied structurally, little attention has been given to the RNCCs that represent the functionally active state of the ribosome. This is in part due to the lack of a simple and reliable procedure for the large-scale preparation of peptidyl-tRNAs. Here we report a new chemoenzymatic approach based on native chemical ligation reaction for the facile synthesis of stably linked peptidyl-tRNAs that were used to determine several structures of RNCCs in the functional pre-attack state of the peptidyl transferase center (PTC) at the highest resolution available to date. These structures reveal a previously unknown role of the ribosome in stabilization of the growing polypeptide within the PTC and suggest an extended entropic trap model that mechanistically rationalizes how ribosome acts with comparable efficiencies upon a multitude of possible growing peptides having various sequences. Our structures also provide new insights into the mechanism of PTC functioning and explain what makes ribosome a versatile catalyst.

## MAIN TEXT

Protein biosynthesis, also known as translation, is a key step in the gene expression pathway, which is catalyzed by the ribosome – one of the most conserved and sophisticated molecular machines of the cell. The ribosome provides a platform for binding the messenger RNA (mRNA) and transfer RNAs (tRNAs), which serve as adaptor molecules and have two functional ends, one carrying the amino acid and the other end containing the anticodon that recognizes the mRNA codon. tRNAs bind to the ribosome in three places: A (aminoacyl), P (peptidyl), and E (exit) sites. The A site binds the incoming aminoacyl-tRNA (aa-tRNA), the P site retains the peptidyl-tRNA carrying the nascent polypeptide chain, and the E site binds deacylated tRNA before it dissociates from the ribosome. The ribosome is composed of two unequal subunits, small and large (30S and 50S in bacteria), which join together to form functional 70S ribosomes. The small subunit decodes genetic information delivered by mRNA, whereas the large subunit covalently links amino acids into a nascent protein, which is then threaded through the nascent peptide exit tunnel (NPET) that spans the body of the large subunit.

Although NPET was initially thought to be a passive conduit for the growing polypeptide chain, it became clear in recent years that it plays an active role in co-translational protein folding (*1*) as well as in the regulation of protein synthesis (*2*). For example, some of the most widely used classes of antibiotics, such as macrolides, bind in the NPET of the bacterial ribosome and interfere with the progression of some growing polypeptides through this tunnel (*3*–*5*). There are also many examples of small molecules and/or metabolites that modulate the rate of translation by binding in the NPET (*2*, *6*). The binding of an antibiotic or a small-molecule modulator to the NPET and the presence of a nascent peptide with a particular sequence results in a slowdown or even a complete arrest of such ribosome-nascent chain complexes (RNCCs) on the mRNA. This happens because the growing peptide adopts a particular conformation inside the NPET and establishes direct contacts with the walls of the tunnel and a small molecule present there at the same time (*2*). Moreover, the synthesis of some nascent chain sequences is intrinsically problematic for the ribosome, with polyproline stretches being the most striking example (*7*).

Although molecular mechanisms underlying nascent chain-mediated ribosome stalling have attracted increasingly great attention in recent years, such events are rare and can occur with relatively few peptide sequences. In fact, evolution has built ribosomes to be able to synthesize most of the cellular proteins without the need for auxiliary factors. Furthermore, ribosome remains an effective catalyst despite the existing chemical diversity of its multiple amino acid substrates as well as possible heterogeneity of growing peptide chain conformations in the NPET. Structural studies of arrested RNCCs can explain why ribosomes become inactive while trying to synthesize particularly problematic amino acid sequences (*8*–*13*). However, the complementary question – “What makes the ribosome a versatile catalyst capable of acting with comparable efficiency upon twenty different aa-tRNA substrates and plenty of possible nascent chain folds within the tunnel?” – always stayed beyond the scope of such studies and could be tackled by the structural analysis of non-arrested RNCCs or their mimics that represent a functionally active state of the ribosome.

In nearly all structural and biochemical studies of arrested RNCCs (*8*–*18*) peptidyl-tRNAs were prepared *in cis* by exploiting the natural peptidyl transferase activity of the ribosome to generate the peptides attached to the tRNAs in the P site. While certainly producing close-to-natural ribosome complexes, this approach allows preparation of homogenous samples of only the peptide-arrested RNCCs, which represent an inactive state of the PTC by definition. Inability to perform pairwise comparisons of structures of arrested vs. non-arrested RNCCs (e.g., with *vs*.without antibiotic or a small molecule; WT stalling motifs *vs*. mutants) hampers our profound understanding of the underlying ribosome stalling mechanisms. Capturing non-arrested nascent peptide chains in the pre-peptidyl transfer state remains challenging and requires either a kinetic control (*19*) or usage of hydrolysis-resistant aa-tRNAs (*20*) to prevent transpeptidation reaction. To date, the only reported structures of the ribosome in the pre-peptidyl transfer state of the PTC contain a minimal possible P-site substrate, fMet-tRNA_i_^Met^, which allows visualizing only a single first amino acid of the nascent peptide but not a longer polypeptide chain within the ribosome tunnel (*19*, *20*). There are no available structures of the ribosome functional complexes containing both aa-tRNA in the A site and peptidyl-tRNA in the P site in the pre-peptidyl transfer state, which is in part due to the lack of a simple and reliable procedure for the large-scale preparation of peptidyl-tRNAs.

Alternatively, peptidyl-tRNAs could be prepared using a combination of synthetic and biochemical techniques and subsequently added *in trans* to ribosomes in order to form the desired RNCCs (*21*). This approach is based on the *in vitro* translation-independent preparation of the peptidyl-tRNAs, allows incorporation of unnatural amino acids, fluorescent or stable isotope labels for FRET and NMR studies, and, in general, provides more flexibility and control over what is being charged to the tRNAs. In order to be more suitable for structural studies, especially for X-ray crystallography or NMR, the peptide moiety of the peptidyl-tRNA must be amide-linked to the CCA-end of the tRNA to prevent spontaneous hydrolysis and deacylation during the timecourse of experiments (*18*, *20*). Although such hydrolysis-resistant tRNAs do not represent natural substrates *per se*, they have not only been shown to be structurally indistinguishable from native tRNA substrates (*19*) but are also active in transpeptidation when placed in the A site and combined with native aa-tRNA in the P site (*20*, *22*), and, therefore, represent a reasonable approximation of the reactive state, which is suitable enough to allow mechanistic hypotheses to be formulated.

Here we report a new chemoenzymatic approach based on native chemical ligation reaction for the facile synthesis of stably linked peptidyl-tRNAs carrying nascent peptide chains of the desired sequences that can be used in a wide range of structural and/or biochemical studies of both arrested and non-arrested RNCCs. We also report several structures of non-arrested RNCCs in the pre-attack state of the peptidyl transferase center (PTC) at the highest resolution available to date. These structures reveal a previously unknown role of the ribosome in stabilization of the growing polypeptide chain within the PTC and suggest an extended entropic trap model that mechanistically rationalizes how ribosome acts with comparable efficiency upon a multitude of possible nascent chain sequences. Moreover, unlike previous views that ribosome generates the α-helical conformation at the C-terminus of the nascent peptide, detailed analysis of our structures suggests that, regardless of the sequence, the C-terminal non-proline residues of the nascent peptide emerge in the uniform zigzag β-strand-like conformation that is stabilized by the intricate network of H-bonds provided by the 23S rRNA nucleotides.

## RESULTS

### NCL-based synthesis of non-hydrolyzable peptidyl-tRNAs

The synthesis of non-hydrolyzable peptidyl-tRNAs is challenging because there are no enzymes that can directly attach desired peptides to a deacylated tRNA. So far, only one method for the preparation of non-hydrolyzable peptidyl-tRNA was established involving customized solid-phase synthesis of the precursors (*23*) that can be barely performed in a standard molecular biology laboratory precluding its wide use by researchers. Moreover, this approach requires laborious tRNA engineering with DNA enzymes as well as enzymatic ligation steps to include the native tRNA nucleoside modifications (*23*). Thus, the utility of the *in trans-RNCC* reconstitution approach is greatly limited by the lack of a facile and reliable procedure for the large-scale synthesis of stable peptidyl-tRNAs.

To bridge this gap, we have established a new simple method for the semi-synthesis of non-hydrolyzable peptidyl-tRNAs using the native chemical ligation (NCL) reaction, which was originally developed for linking unprotected peptide fragments with each other under mild reaction conditions (*24*). NCL is based on a reaction between an activated C-terminal thioester of the first peptide fragment and the N-terminal cysteine residue of the second peptide fragment resulting in the formation of a native peptide bond between the two fragments. In the method that we report here, we used aminoacyl-tRNA charged with cysteine as the C-terminal reactant, while as the N-terminal reactant, we used a peptide with the desired sequence and activated with the thiobenzyl (TBZ) group, which is now available commercially as a C-terminal modification through the majority of vendors providing peptide synthesis. Previously, the Micura group utilized NCL for the synthesis of short peptidyl-tRNA mimics comprising only the CCA-ends (*25*). However, those early procedures relied heavily on laborious and advanced organic synthesis methods to prepare the non-standard solid supports, essentially making them inaccessible to a general molecular biology laboratory. Moreover, flexizymes have been used in the past to charge 3’-NH2-tailed tRNAs with a single amino acid (*26*), which in principle should work for peptides as well. However, to the best of our knowledge, successful applications of the flexizyme-based methodology for charging amino-tailed tRNAs with a peptide have not been reported to date. The advantage of our chemoenzymatic approach is that it utilizes only commonly available equipment, affordable chemicals, and universally available commercial peptide synthesis services and, therefore, could be employed virtually by any laboratory.

The overall procedure includes three steps (**Figure 1A**): (i) tRNA-tailing to replace the 3’-terminal regular adenosine-3’-OH of the CCA-end with its amino-substituted adenosine-3’-NH_2_ analog (*27*, *28*); (ii) enzymatic charging of the tailed 3’-NH2-tRNA with cysteine by the aminoacyl-tRNA-synthetase (*27*, *28*); and (iii) native chemical ligation of the TBZ-activated peptide with cysteinyl-tRNA to yield the final product. It would have been logical to use cysteine-specific tRNA^Cys^ to prepare non-hydrolyzable cysteinyl-NH-tRNA needed for the NCL. However, in our pilot experiments, we have found that methionine-specific aminoacyl-tRNA-synthetase (MetRS) can efficiently mischarge the tailed initiator 3’-NH_2_-tRNA_i_^Met^ with cysteine resulting in >90% yields (**Figure 1B**, lane 2 *vs*. 3), which likely happens due to the compromised editing function of this ARSase as a result of its inability to hydrolyze non-hydrolyzable aa-tRNA (*29*). The main advantage of using the initiator tRNA_i_^Met^ from *E. coli* instead of tRNA^Cys^ as the P-site substrate for the subsequent ribosome structural studies stems from its high affinity for the ribosomal P site, ensuring the proper binding of the tRNA body to the ribosome. Moreover, unlike tRNA^Cys^, the procedure for large-scale preparation of tRNA_i_^Met^ is well established (*20*, *27*, *28*, *30*), and this tRNA is easy to purify and is readily available. Alternatively, cysteine-specific tRNA^Cys^ in combination with the corresponding cysteinyl-tRNA-synthetase can be used at this step. The apparent limitation of our procedure is that the ultimate (C-terminal) amino acid of the peptide chain must always be Cys, which is dictated by the chemistry of NCL reaction. However, the same limitation also exists with the widely used peptide/protein-only NCL, where multiple approaches have been utilized to overcome this limitation and extend the NCL reaction to other amino acids (*31*). In principle, the same strategies could be used to extend our methodology beyond Cys in the last position of the peptide.

**Figure 1.**
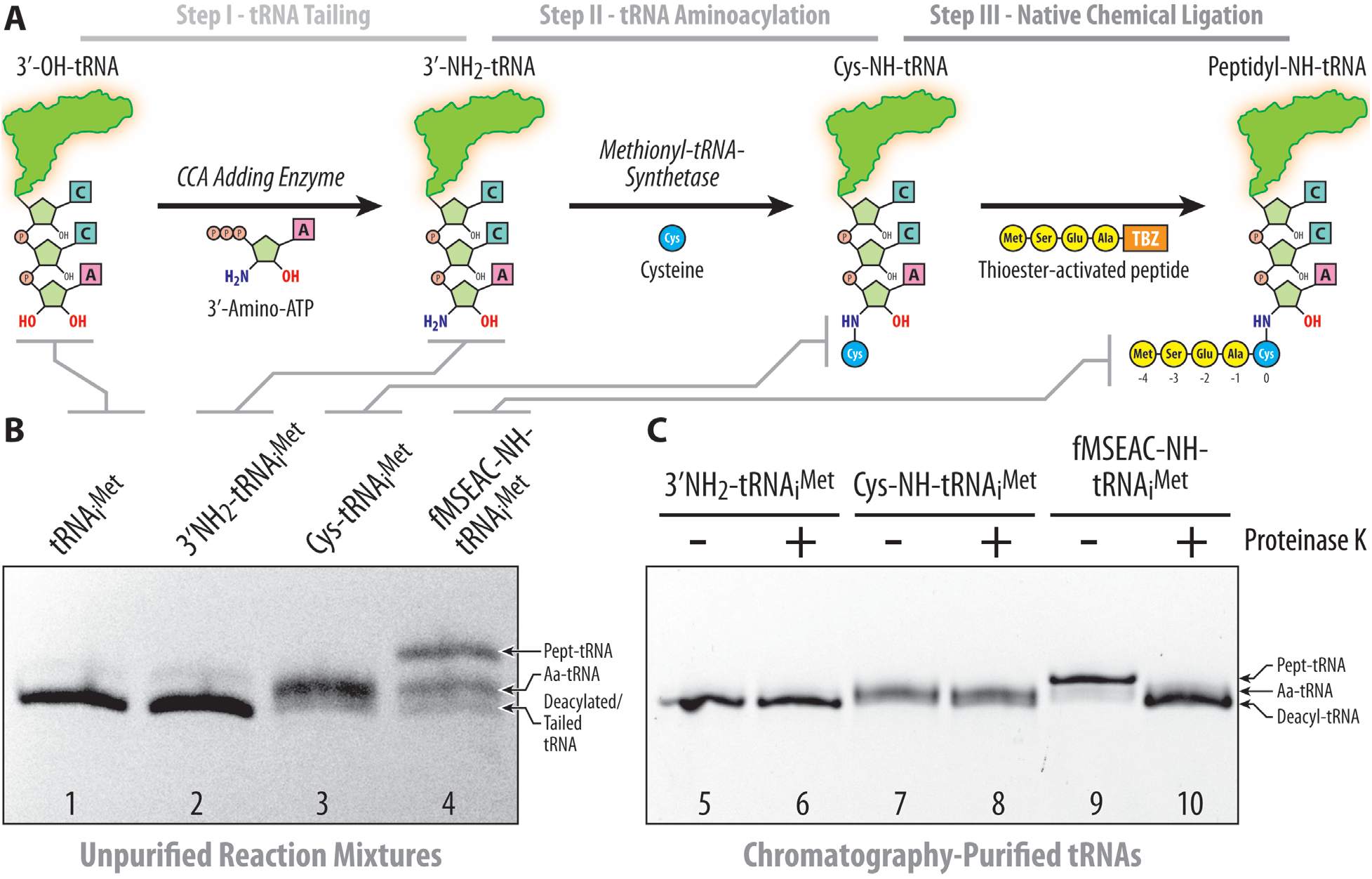
Synthesis of full-length non-hydrolyzable peptidyl-tRNAs by native chemical ligation. (**A**) Steps in preparation of non-hydrolyzable peptidyl-tRNAs. The first tRNA tailing step was performed via the exchange of natural 3’-OH-AMP moiety of tRNA with the 3’-NH_2_-AMP by using the CCA-adding enzyme (*27*). (**B, C**) Electrophoretic separation of native *E. coli* tRNA_i_^Met^ (lane 1) after (i) tRNA tailing reaction (lanes 2, 5, 6); (ii) MetRS-driven aminoacylation with cysteine (lanes 3, 7, 8); (iii) native chemical ligation reaction (lanes 4, 9, 10). In panel (c), samples were treated (+) or not treated (-) with proteinase K. Note the slower mobility of tRNA after the NCL (lane 3 *vs*. 4) and its sensitivity to the proteinase K treatment (lane 9 *vs*. 10). Electrophoresis of the tRNA samples was performed in 20-cm long 8% PAAG with 7 M urea and stained with ethidium bromide.

For the first NCL trials, we have selected several model peptides of different lengths and sequences that would also be interesting to study beyond the scope of this work (**Supplementary Figure 1A**). We should note that some of the peptide sequences in our list represent known ribosome stalling motifs, which, however, cause translation arrest only in the presence of a particular drug (such as PTC-targeting antibiotic chloramphenicol in the case of *hns*, or macrolide antibiotics in the case of *ermDL* gene products), but otherwise are translated normally and should be considered as non-arresting peptides in the absence of the corresponding drug molecules, thereby representing merely model cases for our structural studies. Using the TBZ-activated peptides (purchased from NovoPro Biosciences, China) and the Cys-NH-tRNA_i_^Met^ as the N- and C-terminal fragments for the NCL reaction, respectively, we show that most of the selected peptides can be efficiently ligated to the Cys-NH-tRNA_i_^Met^ with yields exceeding 50% (**Figure 1B**, lane 3 *vs*. 4; **Supplementary Figure 1B**). We used treatment with proteinase K, which hydrolyzes the amide bond between the peptide moiety and the tRNA (*32*), to confirm the presence of the peptidyl moiety attached to the tRNA_i_^Met^ (**Figure 1C**, odd *vs*. even lanes). Even by using the previously reported conditions for peptide ligation (*24*), the efficiency of NCL was already sufficiently high (>50%). Nevertheless, by searching for optimal reactant concentrations, the types and combinations of catalysts and denaturing agents used for the NCL reaction (see **Methods**), we have achieved >70% reaction efficiency for several peptides. Thus, by using the NCL approach, we have synthesized a set of non-hydrolyzable full-length peptidyl-tRNAs carrying various peptide sequences at their CCA-ends, which were then purified by reverse-phase chromatography and used directly for ribosome complex formation and structure determination. We have also used the same NCL-based procedure, but skipping the tRNA-tailing step, to prepare regular ester-linked MSEAC-O-tRNA^Cys^. The yield of this reaction was comparable with the yields of non-hydrolyzable peptidyl-tRNAs, suggesting that our method is not limited to the preparation of only amide-linked peptidyl-tRNAs.

### In trans addition of the peptidyl-tRNAs results in the formation of functionally relevant pre-peptidyl transfer ribosome complexes

Next, to check whether the obtained full-length peptidyl-tRNAs can be used for the structural studies of the non-arrested RNCCs, we prepared a complex of *Thermus thermophilus* 70S ribosome containing fMSEAC-NH-tRNA_i_^Met^ in the P site as well as a full-length aa-tRNA (Phe-NH-tRNA^Phe^) in the A site, crystallized it, and determined its structure at 2.4Å resolution (**Figure 2A, B**). The observed high-quality electron density maps for both A- and P-site tRNA substrates (**Figure 2C**) allowed unambiguous modeling of the four out of five amino acid residues in the fMSEAC peptide moiety attached to the CCA-end of the P-site tRNA (**Figure 2D-F**). We observed no electron density for the N-terminal formyl-methionine residue of the fMSEAC peptide, which is likely due to the lack of its coordination with the surrounding nucleotides of the 23S rRNA resulting in its high flexibility.

**Figure 2.**
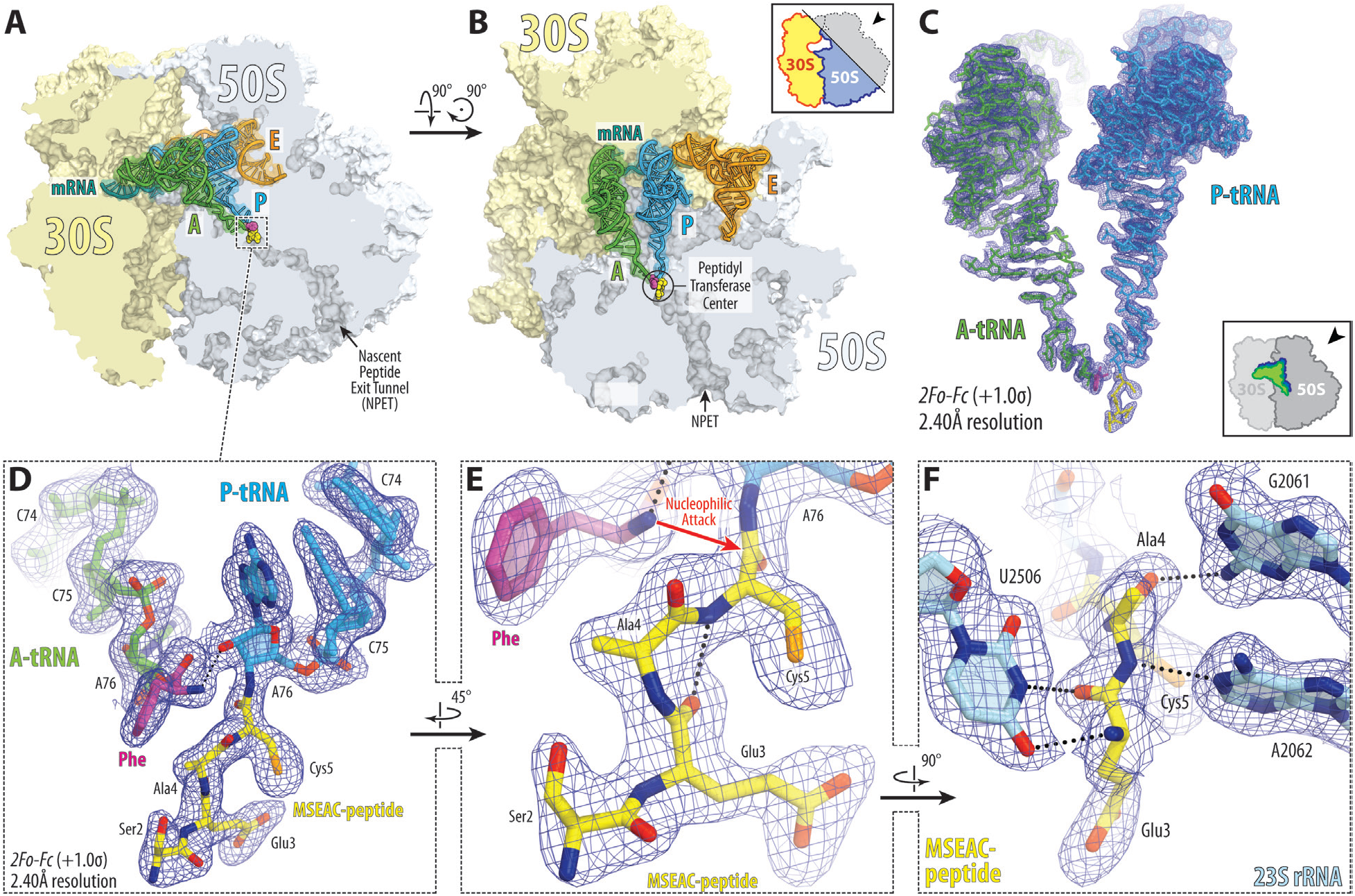
Structure of the 70S ribosome in complex with fMSEAC-peptidyl-tRNA. (**A, B**) Overviews of the *T. thermophilus* 70S ribosome containing full-length aa-tRNA (Phe-NH-tRNA^Phe^) and peptidyl-tRNA (fMSEAC-NH-tRNA_i_^Met^) in the A and P sites, respectively, viewed as cross-cut sections through the ribosome. The 30S subunit is shown in light yellow, the 50S subunit is in light blue, the mRNA is in teal, and the A-, P-, and E-site tRNAs are colored green, blue, and orange, respectively. The phenylalanyl and peptidyl moieties of the A- and P-site tRNAs are colored magenta and yellow, respectively. (**C**) High-resolution (2.4Å) *2F_o_-F_c_* electron difference Fourier map (blue mesh) of the ribosome-bound A- and P-site tRNAs. The refined models of tRNAs are displayed in their respective electron density map contoured at 1.0σ. The entire bodies of the A- and P-site tRNAs are viewed from the back of the 50S subunit, as indicated by the inset. Ribosome subunits are omitted for clarity. (**D-F**) Close-up views of the CCA-ends of the A- and P-site tRNAs carrying phenylalanyl (magenta) and fMSEAC-peptidyl (yellow) moieties, respectively. Nitrogens are colored blue; oxygens are red. H-bonds are shown by black dotted lines. Note that the H-bond between the α-amino group and the 2’-OH of the A76 of the P-site tRNA is pivotal for optimal orientation of α-amine for an in-line nucleophilic attack onto the carbonyl carbon of the P-site substrate. Interactions of the MSEAC-peptide with the nucleotides of the 23S rRNA (light blue) are shown in (f).

Next, we aligned our new structure containing fMSEAC-peptidyl-tRNA with the previously published structures of the 70S ribosome in the pre-attack state containing either the non-hydrolyzable amide-linked (**Supplementary Figure 2A**) (*20*, *28*) or native ester-linked full-length aa-tRNAs in the A and P sites (**Supplementary Figure 2B**) (*19*). We found no major structural differences in the positions of the CCA moieties (**Supplementary Figure 2A, B**) or any of the key 23S rRNA nucleotides around the PTC (**Supplementary Figure 2C, D**), suggesting that, despite the peptidyl-tRNA being added to the ribosome *in trans*, our new structure represents a functionally relevant pre-attack state of the peptidyl-tRNA (**Figure 2E**). In particular, the orientation of the attacking α-amino group of the aa-tRNA relative to the carbonyl carbon of the P-site substrate is nearly identical between the structures harboring amide-linked *vs*. native ester-linked full-length aminoacyl- and peptidyl-tRNAs in the A and P sites, respectively (**Figure 2E; Supplementary Figure 2A, B**) (*20*), indicating that stable amide-linked aminoacyl- and peptidyl-tRNAs are adequate mimics of their native labile ester-linked counterparts.

It is important to emphasize that using stable aminoacyl-/peptidyl-tRNAs allows trapping ribosome in the pre-peptidyl-transfer state because PTC is unable to cleave the amide bond connecting peptide to the P-site tRNA so that peptidyl-tRNA is unable to donate its fMSEAC-peptide moiety to form a peptide bond with the A-site amino acid residue. This methodology of RNCC preparation is principally different from previous approaches used to produce stalled RNCCs, in which translation arrest happens not because substrates are chemically unreactive but because either the key nucleotides of the catalytic site are perturbed (*9*, *33*), or incoming aa-tRNA cannot efficiently accommodate into the A site (*11*), or the C-terminal part of the peptidyl-tRNA adopts non-productive conformation. From this perspective, despite featuring unreactive substrate analogs in the PTC, our ribosome complexes represent non-arrested RNCCs because we observe not only the full accommodation of the A-site tRNA (**Supplementary Figure 2A, B**) but also no major perturbations of the key PTC nucleotides (**Supplementary Figure 2C, D**). Thus, this is the first structure of the non-arrested RNCC containing full-length aminoacyl- and peptidyl-tRNAs that also provides the highest-resolution view of the PTC in the functional pre-attack state immediately before peptide bond formation (**Figure 2E**).

### An extensive H-bond network tightly coordinates growing peptide chains

The structure of fMSEAC-peptidyl-tRNA in the ribosomal NPET reveals tight coordination of the PTC-proximal part of the nascent polypeptide by the elements of the ribosome. In particular, we observe four new hydrogen bonds (H-bonds) between the nucleotides of the 23S rRNA and the three C-terminal amino acid residues of the nascent peptide, as well as an additional intramolecular H-bond between the C-terminal residues of the nascent peptide. The carbonyl and amide groups of the penultimate (−1) residue in the fMSEAC peptide (Ala4) are coordinated by the universally conserved 23S rRNA nucleotides G2061 and A2062, respectively (**Figures 2F; 3A, B**, HB-1 and HB-2). At the same time, the N3 and O4 atoms of the nucleotide U2506 form two additional H-bonds with the carbonyl and amide groups of the (−2) residue of the fMSEAC peptide (Glu3), respectively (**Figures 2F; 3A, B**, HB-3 and HB-4). Another interesting intramolecular H-bond is formed between the amide group of the ultimate residue (Cys) of the peptide and the carbonyl group of the (−2) amino acid (Glu3) (**Figures 2E; 3A**, HB-5). Consistent with previous structural studies, the carbonyl group of the last amino acid residue of the peptide is coordinated by the nucleotide A2602 of the 23S rRNA via a water molecule (referred to as W2 in ref. (*20*)) that was previously proposed to play a key role in stabilizing the tetrahedral oxyanion intermediate of the transition state (**Supplementary Figure 3**) (*20*, *34*). We also observe the two other tightly coordinated water molecules in the PTC (referred to as W1 and W3 in ref. (*20*)) that were suggested to take parts in the formation of an intricate network of H-bonds, the proton wire, needed for efficient deprotonation of the attacking α-amine during the initial rate-limiting step and the subsequent breakdown of the transition-state intermediate (**Supplementary Figure 3**) (*20*).

**Figure 3.**
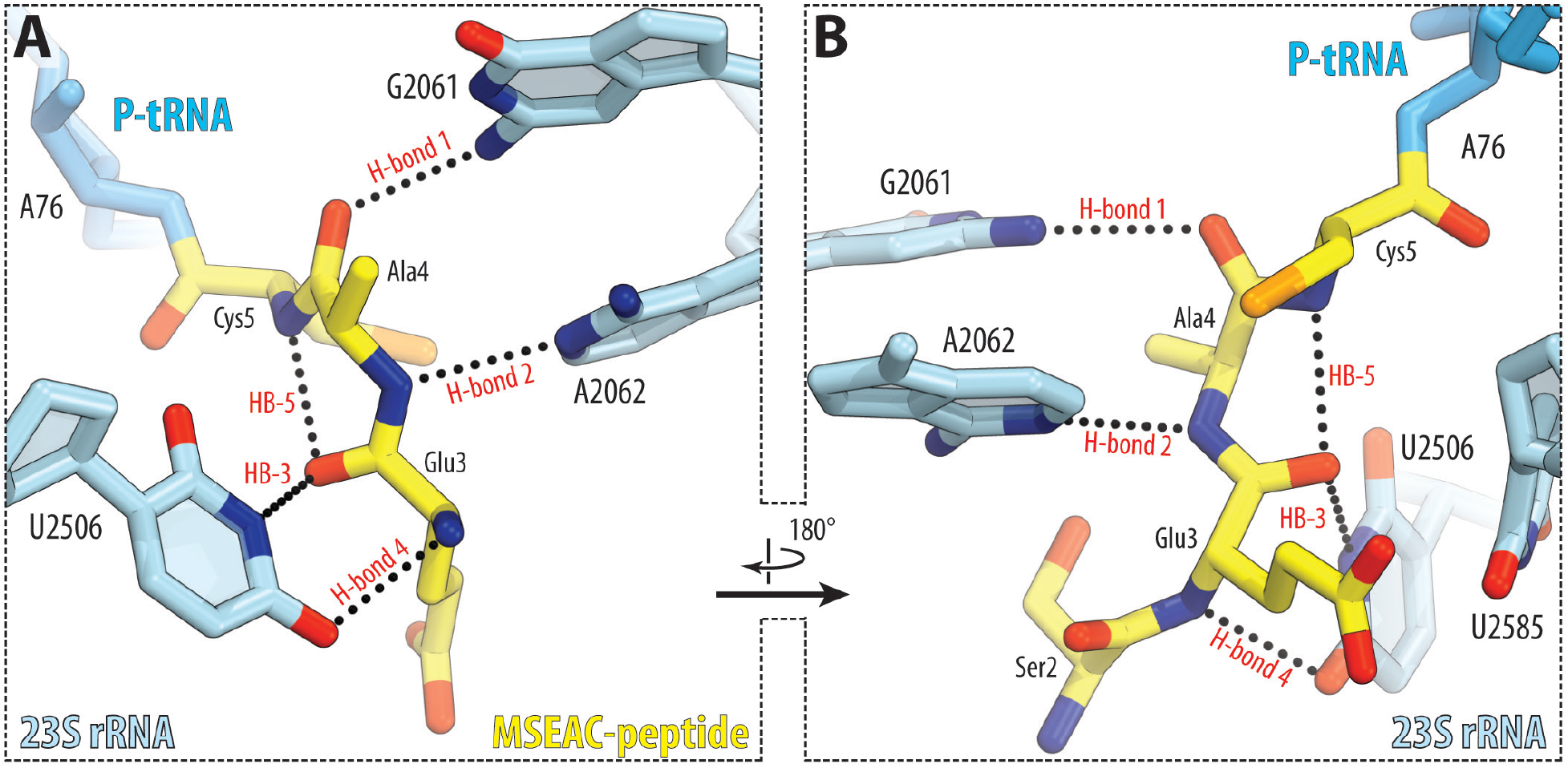
Coordination of the PTC-proximal part of the nascent peptide in the NPET. (**A, B**) Close-up views of the interactions between the ribosome-bound fMSEAC-peptidyl-tRNA (yellow) and the nucleotides of the 23S rRNA (light blue). H-bonds are shown by black dotted lines and are annotated as HB-1 through 5. Nitrogen atoms are shown in blue, and oxygens are red.

Ribosome catalyzes peptide bond formation by (i) stabilizing the transition-state intermediate and (ii) ensuring that the nucleophilic attack by the α-amino group of the aa-tRNA is coordinated with its efficient deprotonation during the rate-limiting step of the reaction (*20*, *35*, *36*). However, for the reaction to even start, the two reactants must be optimally positioned relative to each other in the PTC (*37*). The observed here tight coordination of the C-terminus of the growing peptide inside the NPET might not only (i) ensure the proper fixation of the carbonyl group of the peptidyl-tRNA in the PTC required for efficient attack by the nucleophile to occur but also (ii) prevent premature peptidyl-tRNA drop-off while the peptide remains relatively short (*38*, *39*). Although the contribution of the ribosome to the A- and P-site substrate stabilization was evident from the previous high-resolution structures (*20*, *28*, *34*), it was not known that ribosome stabilizes more than just the carbonyl group of the C-terminal residue of the growing peptide. While universally conserved and essential nucleotides A2451, A2602, and C2063 of the 23S rRNA have been suggested to participate in deprotonation of the attacking α-amino group (*20*), the individual roles of other important PTC nucleotides remained unclear. The observed here network of H-bonds provided by the universally conserved (G2061, A2062, U2506) and also functionally essential (G2061, U2506) nucleotides of the PTC (*40*–*42*) suggests their primary role in peptidyl-tRNA substrate stabilization. It is also evident from the structure that other nucleobases at positions 2061, 2062, and 2506 would be unable to form the same set of H-bonds rationalizing their evolutionary conservation. Curiously, the number of observed H-bonds established by the ribosome with the C-terminal part of the growing peptide is in striking contrast to the lack of any visible H-bonds with the peptide’s N-terminal residues, suggesting that regardless of the length, only the C-terminal residues of the growing peptide need to be coordinated for efficient protein synthesis.

### Nascent peptides emerge in the uniform β-strand-like conformation at the PTC

In contrast to many published cryo-EM structures of various RNCCs, most of which represent arrested (and thus inactive) ribosome complexes, in which PTC is unable to catalyze transpeptidation, our structure provides the first snapshot of the PTC in the functional pre-attack state with both A- and P-site substrates bound. Nevertheless, comparisons of our structure with the available highest-resolution cryo-EM structures of several arrest peptides revealed that the overall trajectories of the C-terminal segments of SpeFL (2.7Å) (*12*), ErmDL (2.9Å) (*11*), or VemP (2.9Å) (*13*) peptides in the NPET are similar to the path of the fMSEAC peptide (**Supplementary Figure 4A-C**) (*11*–*13*). The differences in the paths of the peptides become more prominent as we move further away from the C-terminus. The highest similarity in the peptide trajectories is observed between the structures of the fMSEAC-peptidyl-tRNA and the short tripeptidyl-tRNA analogs carrying MAI, MTI, or MFI tripeptides (*43*) that exhibit nearly identical positions of not only the main-chain but also the Cβ atoms (**Supplementary Figure 4D-F**) (*43*). Because the latter structures also represent non-arrested pre-peptidyl transfer ribosome complexes and also contain peptides with various sequences, it is tempting to propose that any three C-terminal residues (except for prolines) of the growing peptide would always emerge in the same uniform β-strand-like conformation that is strictly enforced by the PTC via multiple H-bonds described above. In principle, there is enough space in the pockets next to the side chains of residues at position −2 (Glu in MSEAC) or position 0 (Cys in MSEAC) to accommodate any of the proteinogenic amino acids. However, the side chain of the residue at position −1 (Ala in MSEAC) points towards the confined space of the ribosomal A site raising the possibility that a large side chain at this position might not be able to fit snugly resulting in sequence-specific alterations of the peptide path in the NPET.

To address this possibility and check whether or not the penultimate residue of the nascent peptide can affect its trajectory in the NPET, we synthesized and purified peptidyl-tRNA carrying model fMRC and fMTHSMRC peptides (**Supplementary Figure 1B**, lanes 2 and 5) with the largest possible amino acid side chain (Arg) in the penultimate position using our NCL-based approach and determined their structures in complex with the ribosome. Except for the C-terminal Leu residue changed to Cys, which does not have any impact on activity (*44*), both of these sequences represent short and long versions of the macrolide-dependent arrest peptide ErmDL, respectively (*11*, *33*, *44*). Appearance of either of these peptide sequences in the NPET results in translation arrest and ribosome stalling, but only in the presence of a macrolide antibiotic and only if combined with lysyl- or arginyl-tRNA in the A site (*44*). Otherwise, these sequences are translated normally and do not cause translation arrest.

Crystals containing *T. thermophilus* 70S ribosome in complex with either the P-site fMRC-NH-tRNA_i_^Met^ or fMTHSMRC-NH-tRNA_i_^Met^ and the A-site Phe-NH-tRNA^Phe^ diffracted to 2.5Å and 2.3Å resolution, respectively (**Supplementary Table 1**). The obtained experimental electron density maps for the corresponding peptidyl-tRNAs allowed unambiguous placement of all of their peptide residues (**Figure 4A, D**), including the N-terminal residues for the longer fMTHSMRC-heptapeptide (**Figure 4B, E**). Most importantly, both peptides appear to be coordinated by the same exact network of H-bonds (**Figure 4C**) as the three C-terminal residues of the fMSEAC-peptide (**Figure 2F**). Moreover, the side chains of penultimate arginines in both peptides establish electrostatic interactions with the phosphate of G2505, further anchoring these peptides in the NPET (**Supplementary Figure 5A, B**). Surprisingly, but despite being longer than fMSEAC, for which only the four C-terminal residues could be resolved, all residues of the fMTHSMRC-heptapeptide are visible in the electron density map. Most likely, this is due to the prominent π-π stacking interaction of the histidine residue with the nucleobase of A2062 resulting in the overall higher rigidity of the entire ribosome-bound peptide (**Supplementary Figure 5C**).

**Figure 4.**
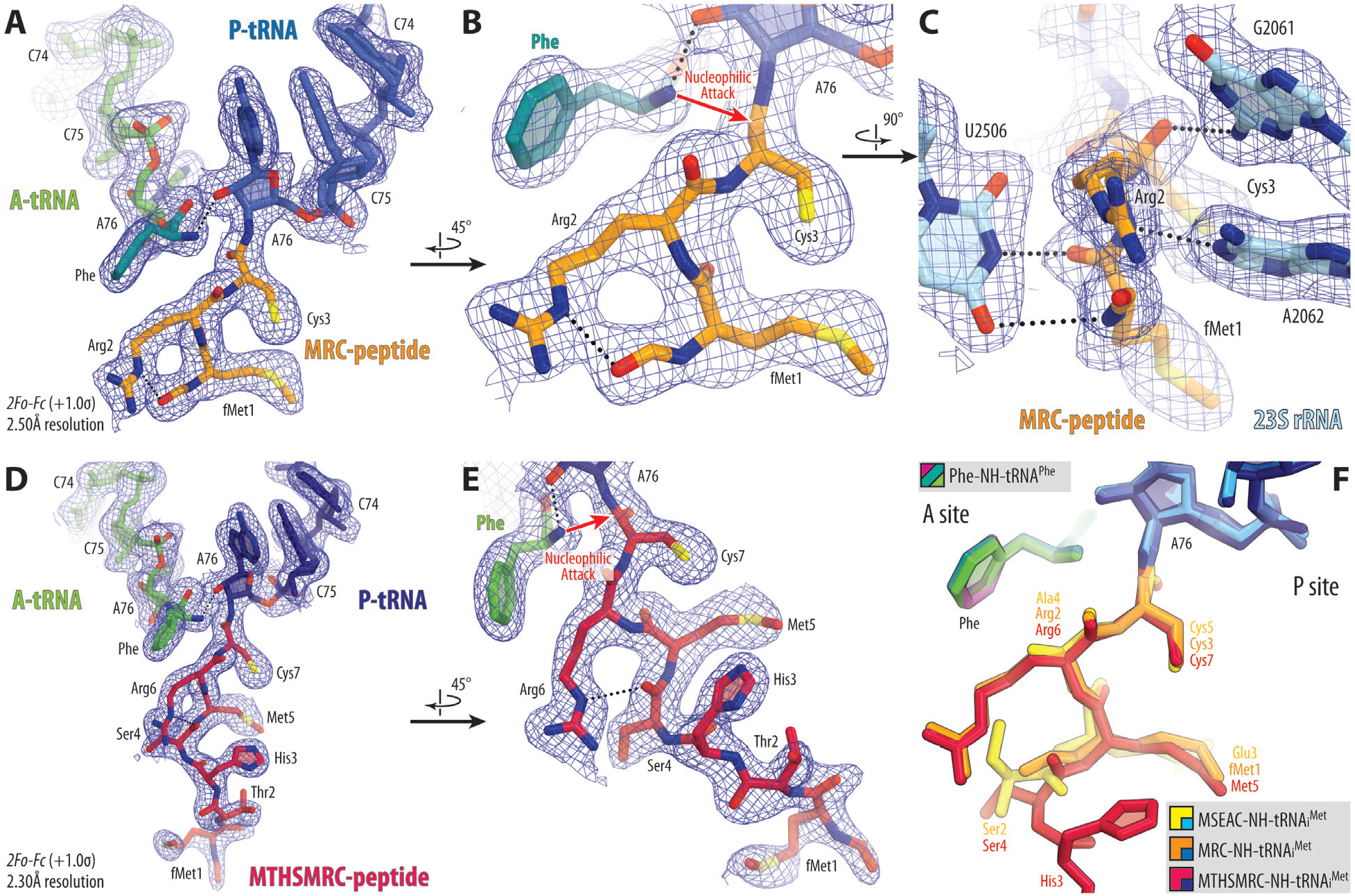
Structure of the 70S ribosome in complex with fMRC- and fMTHSMRC-peptidyl-tRNAs. (**A-E**) Close-up views of the 2*F_o_-F_c_* electron difference Fourier map (blue mesh) of the ribosome complexes containing P-site MRC-NH-tRNA_i_^Met^ (A-C, blue with peptide highlighted in orange) or MTHMRC-NH-tRNA_i_^Met^ (D-E, navy with peptide highlighted in crimson) and A-site Phe-NH-tRNA^Phe^ (green). (**F**) Superpositioning of the current three ribosome structures in complex with the full-length tRNAs carrying either fMSEAC (yellow), or fMRC (orange), or fMTHSMRC (crimson) peptides in the P site. Note that the path of a peptide in the exit tunnel is not affected by the size of amino acid in the penultimate position.

Remarkably, the presence of the bulky arginine side chain in the penultimate position of fMRC or fMTHSMRC peptides does not affect their paths in the NPET (**Figure 4F**), supporting the hypothesis that, regardless of the sequence, the C-terminal non-proline residues of the nascent peptides emerge in the same uniform zigzag β-strand-like conformation that is stabilized by the intricate network of H-bonds provided by the universally conserved nucleotides G2061, A2062, and U2506 of the 23S rRNA. This experiment-driven hypothesis contradicts the previous theoretical calculations-inspired view that ribosome generates the α-helical conformation at the C-end of the nascent peptide (*45*). The observed excessive fixation of the PTC-proximal part of the peptide in the NPET is likely to be important for the efficient catalytic function of the PTC.

## DISCUSSION

The main finding of this study is the intricate network of H-bonds that tightly coordinate the C-terminal part of the nascent peptide in the PTC of the ribosome. All of these interactions involve only the main-chain H-bond donor and acceptor atoms of the nascent polypeptide and, therefore, are sequence-independent that is critical for the ribosome’s ability to translate virtually any sequences. While the sole existence of these H-bonds in the PTC does not necessarily mean that they are required for efficient peptide bond formation, there are several studies that speak in support of this hypothesis. For example, it has been demonstrated that the reactivity of a peptidyl-tRNA is modulated by the length of its nascent peptide chain: peptidyl-tRNAs with longer peptide chains react with puromycin (PMN) more rapidly than those with shorter ones (*46*–*48*). Based on this data, it has been hypothesized that the differences in the peptide bond formation rates with peptides of different lengths are due to tighter fixation of the longer nascent chains, and hence, better positioning of the reactive groups in the PTC (*47*). Analysis of our structural data rationalizes this hypothesis. Indeed, while formyl-methionine forms only a single H-bond out of five possible peptide-stabilizing H-bonds, dipeptides and tripeptides can form three and five H-bonds, respectively, apparently being more tightly restrained in the transpeptidation-competent conformation. As evident from our structures, having more than three amino acid residues in a growing peptide chain does not provide any additional stabilization suggesting that the rate of transpeptidation should reach a plateau at 3-aa peptide length, a prediction which is in excellent agreement with the previous biochemical data (*48*).

Interesting is the inability of proline residues to form some of these H-bonds that can rationalize the apparent redundancy of these interactions involving not just one but the three C-terminal residues of the growing peptide (positions 0, −1, and −2). Despite a number of previous structural and functional studies (*7*, *49*, *50*), there is no definitive answer to the fundamental question of why ribosome is unable to polymerize more than two consecutive prolines. In this case, ribosome stalling occurs due to the slow peptide bond formation between the peptidyl-Pro-Pro-tRNA^Pro^ and the Pro-tRNA^Pro^ located in the A site (*7*, *49*). Previous biochemical studies suggested that the poor reactivity of the Pro-containing peptides stems from the steric, rather than chemical, properties of this imino acid (*51*). However, the mechanistic understanding of why peptidyl-Pro-Pro-tRNAs are especially inactive as P-site substrates is still lacking. *In silico* modeling reveals that appearance of a single proline residue at any of the three C-terminal positions of the growing peptide chain results in a loss of some but not all of the peptide-restraining H-bonds (**Supplementary Figure 6**), which provide incomplete yet sufficient stabilization of the conformation of the nucleophile acceptor, the carbonyl group, for the transpeptidation reaction to occur. This finding is corroborated by the biochemical data showing a significantly slower (~700 fold) rate of peptide bond formation with the C-terminal proline of the peptidyl-tRNA (*47*). However, the presence of two consecutive prolines in a growing polypeptide chain results in the loss of at least four out of five H-bonds (**Supplementary Figure 7A**) that most certainly negatively affects the efficiency of transpeptidation reaction because poorly stabilized diprolyl-tRNA substrate with “twitching” carbonyl group is unlikely to be a good donor during the nucleophilic attack. What is more striking is that, unlike any other residue, modeling of the proline in the penultimate position of the growing peptide shows a substantial steric clash with the nucleotide A2062 of the 23S rRNA (**Supplementary Figures 6B, 7A**), which is locked in place through symmetric trans A-A Hoogsteen base pair with the residue A2503 suggesting that, during the passage of proline residues, the growing peptide must deviate to the side to avoid this steric hindrance. Comparison of our model with the available structure of the ribosome-bound diprolyl-tRNA analog confirmed our prediction showing an alternate path of the Pro-Pro-containing peptide in the tunnel (**Supplementary Figure 7B**) (*52*). Nevertheless, having two consecutive prolines in the peptide sequence does not result in a complete ribosome stalling (*7*), most likely because the unstabilized wobbling peptidyl-tRNA could still randomly visit the productive conformation(s), and transpeptidation can still occur, albeit at a much slower rate (*7*). However, ribosome stalls completely when the next incoming amino acid is a yet another proline residue (*7*). Most likely, this happens because the non-optimal position of the attacking A-site nucleophile is now combined with the “twitching” of the poorly stabilized carbonyl group of the diprolyl-tRNA substrate in the P site. Thus, it becomes obvious that elongation factor P, which is recruited to the ribosome every time two or more consecutive prolines need to be polymerized, provides stabilization of the otherwise wobbling diprolyl-tRNA substrate in the P site consistent with its previously proposed role in translation as an entropy-decreasing factor (*7*, *49*, *50*).

The efficient protein synthesis requires tight coordination of A- and P-site substrates during all peptide bond formation events (*37*), including the first one. Analysis of our structure also provides insight into the possible role of formylation of initiator tRNA in bacteria. Superpositioning of the ribosome structure containing formylated initiator fMet-tRNA_i_^Met^ in the P site with the new structure of fMSEAC-peptidyl-tRNA reveals nearly identical positions of the carbon and oxygen atoms in the formyl group and those in the carbonyl group of the penultimate residue in the growing peptide chain (**Supplementary Figure 8**). Similar to the carbonyl group, the formyl group is coordinated by the same nucleotide G2061 of the 23S rRNA and, thus, provides additional stabilization to the entire methionine residue attached to the P-site tRNA (**Supplementary Figure 8**). In other words, during the first round of elongation, when only a single amino acid (and not yet a peptide) is present in the P site, the formyl group mimics a dipeptide and provides added coordination to the P-site substrate that otherwise would be available only to a dipeptidyl-tRNA. Consistent with our observation, the inability of non-formylated methionine residue to form the same H-bond with the G2061 results in a 5-fold decrease of the transpeptidation reaction rate (*53*).

Lack of the machinery for formylation of the initiator tRNA in eukaryotes could have been compensated in the course of evolution by acquiring a dedicated protein factor, such as eukaryotic initiation factor 5A (eIF-5A), that provides added stability to the first methionine residue resulting in a more efficient transpeptidation. Although there is compelling evidence that both eukaryotic/archaeal initiation factor eIF-5A and bacterial elongation factor EF-P are needed primarily for the synthesis of polyproline stretches in proteins (*7*, *54*, *55*) and both carry functionally important hyper-modified lysine residues that encroach upon the P-site substrate in the PTC (*52*, *56*), we cannot exclude the possibility that eIF-5A also contributes to the first peptide bond formation as suggested by early reports (*57*–*59*), and it is only later that its role in elongation has also been revealed (*54*, *55*). These roles are not mutually exclusive, and it is conceivable that while diverging from a common ancestor, eIF-5A retained both of the original functions (facilitation of the first peptide bond formation and facilitation of the synthesis of polyproline tracts), whereas EF-P retained only the second function delegating the first one to the initiator tRNA formylation machinery.

## CONCLUSIONS

In summary, here we present a simple, affordable, and fast NCL-based method for the synthesis of stably linked peptidyl-tRNAs carrying nascent peptide chains of the desired sequences (except for the C-terminal Cys) that can be used in a wide range of structural and/or biochemical studies, especially those that are focused at understanding the role of the nascent peptide sequence in the regulation of protein synthesis in response to various environmental cues, such as small molecules and antibiotics. By providing the first 2.3-2.5Å resolution X-ray crystal structures of the RNCCs featuring hydrolysis-resistant aminoacyl-tRNA in the A site and peptidyl-tRNAs in the P site, we demonstrate that synthetic full-length peptidyl-tRNAs can efficiently be introduced into the ribosome *in trans* and yet represent a functionally significant pre-attack state of the PTC. The availability of such peptidyl-tRNAs opens avenues for the structural studies of RNCCs under both stalling and non-stalling conditions in parallel, including rationalization of the context-specific mechanisms of action of many ribosome-targeting antibiotics such as those that target the PTC (*60*, *61*) or the NPET (*33*). Moreover, the detailed analysis of our structural data suggests an answer to the question of why ribosome is unable to polymerize more than two consecutive proline residues without the need for a dedicated facilitator (such as EF-P or eIF-5A). Furthermore, here we provide important insights into the possible role of formylation of the initiator tRNA in the protein synthesis in bacteria. Finally, we propose a hypothesis that the C-terminal segments of all nascent peptides emerge in the same uniform β-strand-like conformation that is yet to be verified experimentally as more structures of non-arrested RNCCs will become available with the advent of the new method for facile synthesis of full-length non-hydrolyzable peptidyl-tRNAs by native chemical ligation.

## AUTHOR CONTRIBUTIONS

E.A.S. with the help from E.V.A. developed the NCL-based method for the synthesis of non-hydrolyzable full-length peptidyl-tRNAs; E.A.S., E.V.A., and Y.S.P designed and performed X-ray crystallography experiments; Y.S.P. supervised the experiments. All authors interpreted the results. E.A.S., E.V.A., and Y.S.P. wrote the manuscript.

## ACKNOWLEDGMENTS

We thank Dr. Ronald Micura and Dr. Maksim Svetlov for the critical reading of the manuscript and valuable suggestions. We thank the staff at NE-CAT beamlines 24ID-C and 24ID-E for help with data collection and freezing of the crystals, especially Drs. Malcolm Capel, Frank Murphy, Surajit Banerjee, Igor Kourinov, David Neau, Jonathan Schuermann, Narayanasami Sukumar, Anthony Lynch, James Withrow, Kay Perry, Ali Kaya, and Cyndi Salbego.

This work is based upon research conducted at the Northeastern Collaborative Access Team beamlines, which are funded by the National Institute of General Medical Sciences from the National Institutes of Health [P30-GM124165 to NE-CAT]. The Eiger 16M detector on 24-ID-E beamline is funded by an NIH-ORIP HEI grant [S10-OD021527 to NE-CAT]. This research used resources of the Advanced Photon Source, a U.S. Department of Energy (DOE) Office of Science User Facility operated for the DOE Office of Science by Argonne National Laboratory under Contract No. DE-AC02-06CH11357.

This work was supported by the National Institutes of Health [R01-GM132302 and R21-AI37584 to Y.S.P.], the National Science Foundation [MCB-1907273 to Y.S.P.], and the Illinois State startup funds [to Y.S.P.].

## METHODS

### Preparation of peptidyl-tRNAs

All reagents and chemicals were obtained from MilliporeSigma (USA). Wild-type deacylated initiator tRNA_i_^Met^ was overexpressed and purified from *E. coli* as described previously (*30*). tRNA tailing was performed in the conditions that were reported previously (*27*) with minor modifications. Briefly, deacylated tRNA_i_^Met^ (40 μM) was incubated at 37°C for 1 hour in a buffer containing 100 mM Glycine-NaOH pH 9.0, 1 mM DTT, 1 mM pyrophosphate, 1 mM 3’-NH_2_-ATP, and 10 μM of the CCA-adding enzyme from *E. coli*. The reaction was terminated by the addition of EDTA to 20 mM, treated with the phenol-chloroform mixture, and precipitated with ethanol. The resulting RNA pellet was dissolved in 20-40 μL of ammonium acetate (pH 5.5), and the solution was desalted via Sephadex G-25 (MilliporeSigma, USA) spin columns (500 μL media per 20-40 μL solution). For aminoacylation of the tailed 3’-NH_2_-tRNA_i_^Met^ with cysteine, 40 μM tRNA was incubated at 25°C for 80 minutes in a buffer containing 100 mM HEPES-KOH pH 8.2, 20 mM MgCl_2_, 7.5 mM KCl, 1 mM DTT, 10 mM ATP, and 1 mM cysteine together with 1 mg/ml methionine-specific aminoacyl-tRNA-synthetase (MetRS) from *E. coli*. Aminoacylation reaction was terminated by the addition of EDTA to 30 mM concentration and then treated with phenol and precipitated with ethanol. The pellet was dissolved in a buffer containing 20 mM HEPES-KOH pH 8.2 and 4 mM TCEP to a final tRNA concentration of 20 μg/μL.

For native chemical ligation, dry thioester-activated peptides fMSEA-TBZ, fMR-TBZ, or fMTHSMR-TBZ (98% purity) carrying thio-benzyl group as the C-terminal modification (NovoPro Biosciences, China) was dissolved in a buffer containing 1M HEPES-KOH pH 7.4 and 6M guanidine-HCl to obtain 50 mM final concentration. Next, 5 μL of the peptide solution were mixed with 15 μL of 400 mM 4-mercaptophenylacetic acid (MPAA) titrated to pH 7.0 with NaOH, and 20 μL of Cys-NH-tRNA_i_^Met^. Also, TCEP pH 6.8 was added to the reaction mixture to 100 mM final concentration. The NCL reaction mixture was incubated for 16 hours at room temperature. The NCL products were purified by HPLC on a 1.7-mL C4 reversed-phase column (Proteo 300, 100×4.6 mm, Higgins Analytical) using 20-mL 0-60% linear gradient of 40% methanol in a buffer containing 20 mM NH_4_CH_3_COO pH 5.5, 400 mM NaCl, 10 mM MgCl_2_, 1 mM EDTA, 1% methanol, 10 mM β-mercaptoethanol. Before applying onto the C4 column, the NCL mixture underwent a buffer-exchange procedure using Amicon Ultra 10K centrifugal filter units (MilliporeSigma, USA). The fractions from the C4 column that contained the desired peptidyl-tRNAs were pooled, ethanol-precipitated, and dissolved in a buffer containing 10 mM NH4CH3COO pH 5.5, 5 mM DTT to 200 μM final concentration. The aliquots were flash-frozen in liquid nitrogen and stored at −80°C until further use in crystallization experiments.

### Proteinase K treatment and gel electrophoresis of peptidyl-tRNA

For proteinase K treatment, 1 μM of the C4-purified fMSEAC-peptidyl-tRNA was incubated with 0.4 mg/mL proteinase K (MilliporeSigma, USA) in a buffer containing 10 mM Tris-Acetate pH 8.0, 10 mM Mg(CH_3_COO)_2_ and 40 mM NH_4_CH_3_COO for 1 hour at 25°C. The reaction was terminated by the addition of 2 volumes of formamide electrophoresis buffer containing 10%β-mercaptoethanol and heating for 2 minutes at 95°C. For separation of the NCL products, we used denaturing 8% (19:1) PAGE in the presence of 7 M urea (20-cm long, 0.4-mm thick gels). Gels were stained with ethidium bromide and visualized with Alpha Imager (Alpha Innotek, USA).

### X-ray crystallographic structure determination

Wild-type 70S ribosomes from *Thermus thermophilus* (strain HB8) were prepared as described previously (*20*, *62*–*64*). Synthetic mRNA with the sequence 5’-GGC-AAG-GAG-GUA-AAA-**AUG**-**UUC**-UAA-3’ containing Shine-Dalgarno sequence followed by methionine (blue) and phenylalanine (red) codons was obtained from Integrated DNA Technologies (USA). Non-hydrolyzable aminoacylated Phe-NH-tRNA^Phe^ was prepared as described previously (*28*) using optimized 3’-NH_2_-tailing and aminoacylation procedures described above (*27*). Complexes of the wild-type *Thermus thermolilus* 70S ribosomes with mRNA and hydrolysis-resistant aminoacyl (A-site Phe-NH-tRNA^Phe^) and peptidyl (P-site fMSEAC-NH-tRNA_i_^Met^, or fMRC-NH-tRNA_i_^Met^, or fMTHSMRC-NH-tRNA_i_^Met^) tRNAs were formed as described previously for deacylated (*64*) or aminoacylated tRNAs (*20*, *28*). Collection and processing of the X-ray diffraction data, model building, and structure refinement were performed as described in our previous publications (*20*, *28*, *63*, *64*). The statistics of data collection and refinement are compiled in **Supplementary Table 1**. All figures showing atomic models were rendered using PyMol software (www.pymol.org).

## DATA AVAILABILITY STATEMENT

Coordinates and structure factors were deposited in the RCSB Protein Data Bank with accession codes:

- **7XXX** for the *T. thermophilus* 70S ribosome in complex with mRNA, aminoacylated A-site Phe-NH-tRNA^Phe^, peptidyl P-site fMSEAC-NH-tRNA_i_^Met^, and deacylated E-site tRNA^Phe^;
- **7YYY** for the *T. thermophilus* 70S ribosome in complex with mRNA, aminoacylated A-site Phe-NH-tRNA^Phe^, peptidyl P-site fMRC-NH-tRNA_i_^Met^, and deacylated E-site tRNA^Phe^;
- **7ZZZ** for the *T. thermophilus* 70S ribosome in complex with mRNA, aminoacylated A-site Phe-NH-tRNA^Phe^, peptidyl P-site fMTHSMRC-NH-tRNA_i_^Met^, and deacylated E-site tRNA^Phe^.

All previously published structures that were used in this work for model building and structural comparisons were retrieved from the RCSB Protein Data Bank: PDB entries 6XHW, 6WDD, 6TC3, 5NWY, 5DGV.

