## Supplementary Information for "Insights into the ribosome function from the structures of non-arrested ribosome nascent chain complexes"

**This file includes:**

I. Supplementary Table 1;

II. Supplementary Figures 1 to 8 with legends;

III. Supplementary References.

### II. SUPPLEMENTARY TABLES

#### Supplementary Table 1. X-ray data collection and refinement statistics.

| ***Crystals*** | | **70S ribosome complex with A-site Phe-NH-tRNA$^{Phe}$, and P-site fMSEAC-NH-tRNA$_i^{Met}$ PDB entry 7XXX** | **70S ribosome complex with A-site Phe-NH-tRNA$^{Phe}$, and P-site fMRC-NH-tRNA$_i^{Met}$ PDB entry 7YYY** | **70S ribosome complex with A-site Phe-NH-tRNA$^{Phe}$, and P-site MTHSMRC-NH-tRNA$_i^{Met}$ PDB entry 7ZZZ** |
|---|---|---|---|---|
| ***Diffraction data*** | | | | |
| Space Group | | $P2_12_12_1$ | $P2_12_12_1$ | $P2_12_12_1$ |
| Unit Cell Dimensions, Å (a x b x c) | | 210.47 x 451.32 x 627.74 | 210.47 x 451.32 x 627.74 | 209.43 x 447.70 x 618.27 |
| Wavelength, Å | | 0.9791 | 0.9792 | 0.9791 |
| Resolution range (outer shell), Å | | 175-2.40 (2.46-2.40) | 191-2.50 (2.56-2.50) | 187-2.30 (2.36-2.30) |
| I/σI (outer shell) | | 8.96 (1.00) | 7.99 (0.93) | 6.54 (0.83) |
| Resolution at which I/σI=1, Å | | 2.40 | 2.50 | 2.30 |
| Resolution at which I/σI=2, Å | | 2.60 | 2.75 | 2.60 |
| CC(1/2) at which I/σI=1, % | | 16.8 | 19.4 | 12.5 |
| CC(1/2) at which I/σI=2, % | | 39.3 | 50.0 | 45.0 |
| Completeness (outer shell), % | | 98.1 (95.5) | 99.6 (98.6) | 97.5 (94.0) |
| $R_{merge}$ (outer shell)% | | 16.2 (139.5) | 18.7 (150.7) | 14.6 (141.0) |
| No. of crystals used | | 1 | 1 | 1 |
| No. of Reflections Used: | 11,101,606 | 11,647,854 | 14,684,473 | 8,504,865 |
| | 2,249,111 | 2,012,356 | 2,100,555 | 2,467,641 |
| Redundancy (outer shell) | | 4.94 (4.80) | 5.79 (4.88) | 3.44 (3.40) |
| ***Refinement*** | | | | |
| Resolution range of the diffraction data included in the refinement, Å | | 110-2.40 | 128-2.50 | 187-2.30 |
| $R_{work}/R_{free}$, % | | 24.2/29.5 | 22.9/28.4 | 23.3/28.2 |
| *No. of Non-Hydrogen Atoms* | | | | |
| RNA | | 200,247 | 200,247 | 200,247 |
| Protein | | 91,022 | 91,010 | 91,072 |
| Ions (Mg, K, Zn, Fe) | | 2,812 | 2,810 | 2,813 |
| Waters | | 4,316 | 4,315 | 4,335 |
| *Ramachandran Plot* | | | | |
| Favored regions, % | | 90.80 | 90.58 | 91.22 |
| Allowed regions, % | | 8.92 | 9.14 | 8.49 |
| Outliers, % | | 0.28 | 0.28 | 0.29 |
| *Deviations from ideal values (RMSD)* | | | | |
| Bond, Å | | 0.008 | 0.009 | 0.009 |
| Angle, degrees | | 1.409 | 1.546 | 1.411 |
| Chirality | | 0.058 | 0.062 | 0.059 |
| Planarity | | 0.008 | 0.008 | 0.008 |
| Dihedral, degrees | | 17.527 | 17.634 | 17.384 |
| Average B-factor (overall), $Å^2$ | | 50.4 | 49.7 | 54.2 |

#### II. SUPPLEMENTARY FIGURES

**A**

| Source Gene Product | Original Peptide Sequence | Translation Arrest in the Presence of | Synthetic TBZ-activated Peptide for NCL | Final peptidyl-tRNA Product |
|---|---|---|---|---|
| ErmDL | MRL | ERY & R/K in the A site | fMR-TBZ | fMRC-tRNA |
| N/A | MKKK | No Stalling | fMKKK-TBZ | fMKKKC-tRNA |
| N/A | MFLRC | Unknown | fMFLR-TBZ | fMFLRC-tRNA |
| ErmDL | MTHSMRL | ERY & R/K in the A site | fMTHSMR-TBZ | fMTHSMRC-tRNA |
| ErmDL | MTHLMRL | No Stalling | fMTHLMR-TBZ | fMTHLMRC-tRNA |
| H-NS | MSEAL | CHL & not G in the A site | fMSEA-TBZ | fMSEAC-tRNA |
| H-NS | MSEALKILN | CHL & not G in the A site | fMSEALKIL-TBZ | fMSEALKILC-tRNA |
| Apidaecin | MNRPVYIPQPRPPHPRL | RF and deac-tRNA | fMNRPVYIPQPRPPHPR-TBZ | fMNRPVYIPQPRPPHPRC-tRNA |

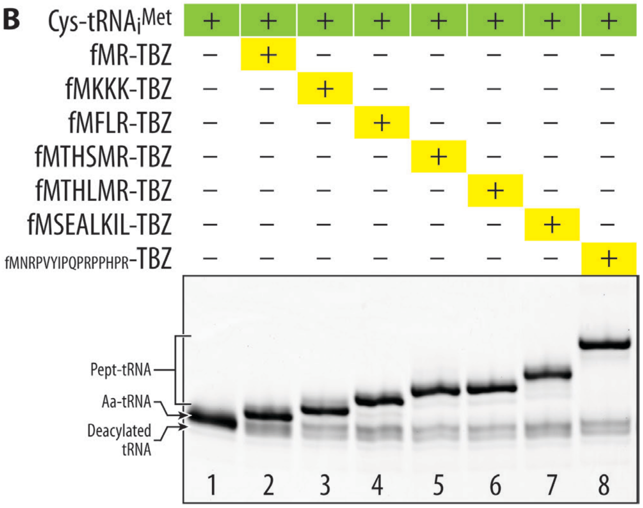


**Supplementary Figure 1. Native chemical ligation of Cys-NH-tRNA$_i^{Met}$ with TBZ-activated peptides of various lengths and sequences.** (**A**) General information about the model peptides chosen for this study: macrolide-sensing ErmDL stalling peptide (*1, 2*); macrolide-resistance peptide MFLRC (*3*); chloramphenicol-sensitive protein H-NS (*4*); proline-rich antimicrobial peptide apidaecin (*5*). ERY, erythromycin or other macrolides; CHL, chloramphenicol; RF, release factor. Peptidyl-tRNA products used in the structural studies are highlighted in grey. (**B**) Electrophoretic separation of crude NCL reaction mixtures. Indicated TBZ-activated peptides and Cys-NH-tRNA$_i^{Met}$ were used as N- and C-terminal reactants, respectively. Electrophoresis was performed in 20-cm long 8% PAAG with 7 M urea and stained with ethidium bromide. Note the slower mobility of tRNA after the NCL that is also proportional to the ligated peptide length (lanes 2-8 *vs.* 1).

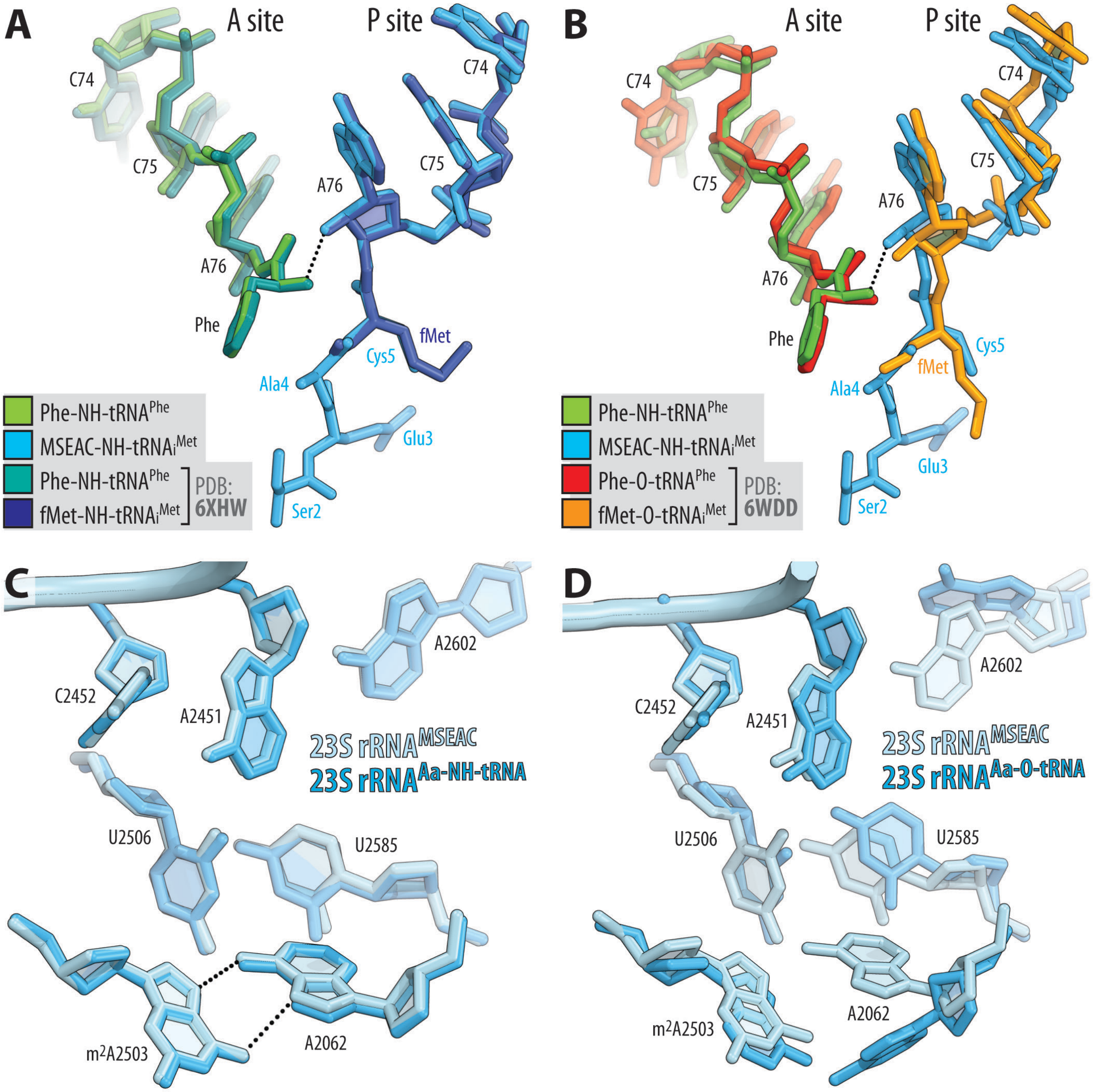


**Supplementary Figure 2. Comparison of the structures of fMSEAC-peptidyl-tRNA with aminoacylated full-length tRNAs.** (**A, B**) Superpositioning of our 70S ribosome structure carrying Phe-NH-$tRNA^{Phe}$ (green) and fMSEAC-NH-$tRNA_i^{Met}$ (blue) in the A and P sites, respectively, with the previously reported structures of ribosome-bound full-length aminoacyl-tRNAs featuring either non-hydrolyzable amide linkages (a, PDB entry 6XHW (*6*)) or native ester bonds (b, PDB entry 6WDD (*7*)) between the amino acid moieties and the ribose of nucleotide A76 of A- and P-site tRNAs. All structures were aligned based on domain V of the 23S rRNA. (**C, D**) Comparisons of the positions of key 23S rRNA nucleotides around the PTC in the same structures. Note that there are no significant differences in the positions of A- or P-site substrates or the PTC nucleotides indicating that the amide-linked aminoacyl and peptidyl-tRNAs represent functionally meaningful analogs of native ester-linked tRNAs.

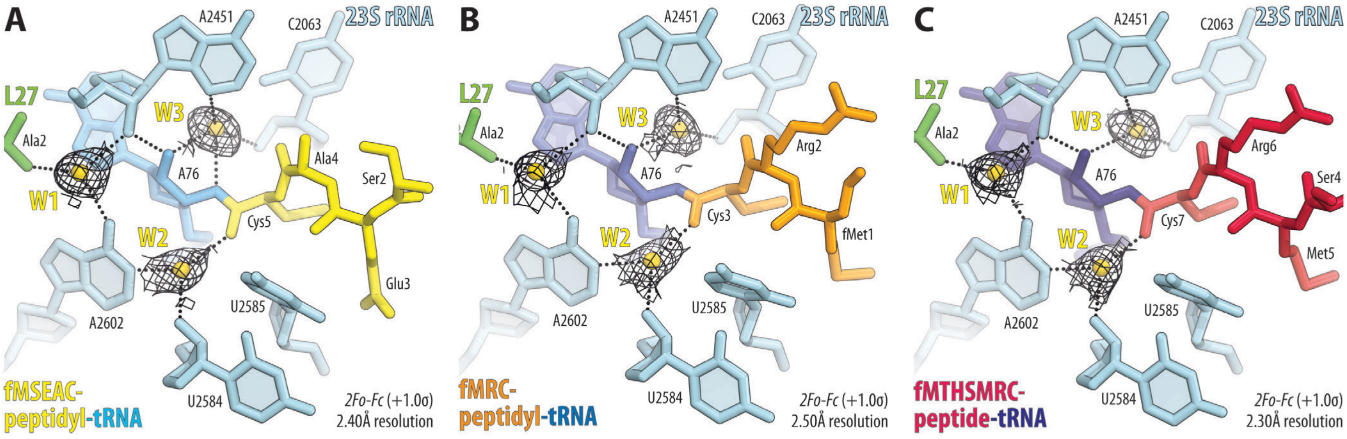


**Supplementary Figure 3. Tightly coordinated water molecules in the pre-attack state of the peptidyl transferase center.** (**A-C**) Close-up views of the 2$F_o$-$F_c$ electron difference Fourier map (black mesh) for water molecules W1, W2, and W3 (yellow; nomenclature from (*8*)) in the pre-peptidyl-transfer complex structures featuring Phe-NH-tRNA$^{Phe}$ in the A site (omitted for clarity) and fMSEAC-NH-tRNA$_i^{Met}$ (A, yellow), fMRC-NH-tRNA$_i^{Met}$ (B, orange), or fMTHSMRC-NH-tRNA$_i^{Met}$ (C, crimson) in the P site. H-bonds are shown by black dotted lines.

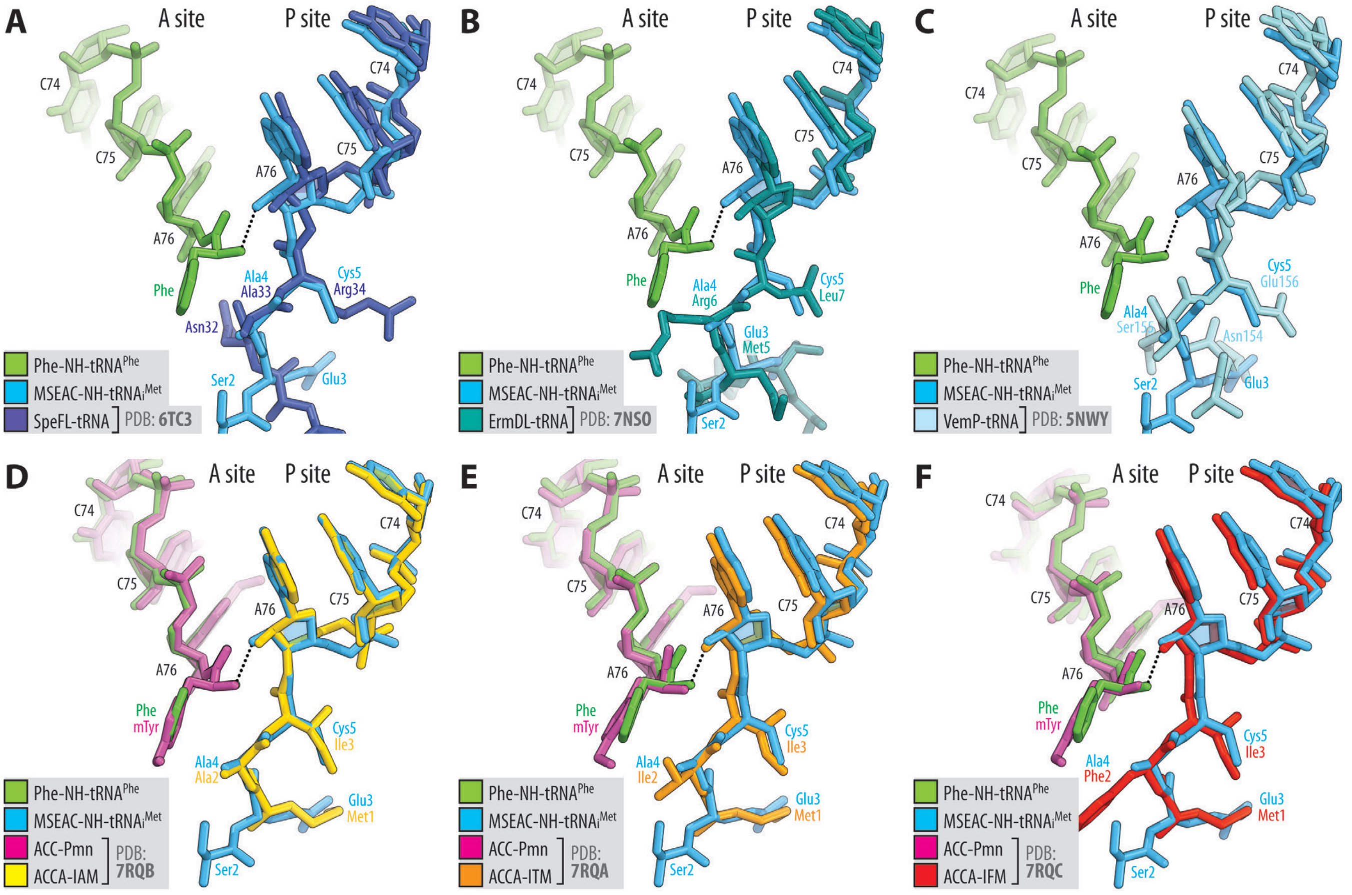


**Supplementary Figure 4. Comparison of the structures of fMSEAC-peptidyl-tRNA with the structures of other ribosome-bound peptidyl-tRNAs.** Superpositioning of the 70S ribosome structure containing A-site Phe-NH-$tRNA^{Phe}$ (green) and P-site fMSEAC-NH-$tRNA_i^{Met}$ (blue) with the previously reported structures of stalled RNCCs carrying full-length peptidyl-tRNAs (**A-C**) or non-stalled RNCCs carrying short non-hydrolyzable tripeptidyl-tRNA analogs (**D-F**). Individual panels show comparisons of the fMSEAC-tripeptidyl-tRNA with the following peptides: (A) SpeFL (dark blue, PDB entry 6TC3 (*9*)); (B) ErmDL (teal, PDB entry 7NSO (*2*)), (C) VemP (cyan, PDB entry 5NWY (*10*)); (D) MAI-tripeptide (yellow, PDB entry 7RQB (*11*)); (E) MTI-tripeptide (orange, PDB entry 7RQA (*11*)), (F) MFI-tripeptide (red, PDB entry 7RQC (*11*)). All structures were aligned based on domain V of the 23S rRNA. Note that the overall path of the MSEAC peptide in our structure is similar to the trajectories of the other peptides in the NPET.

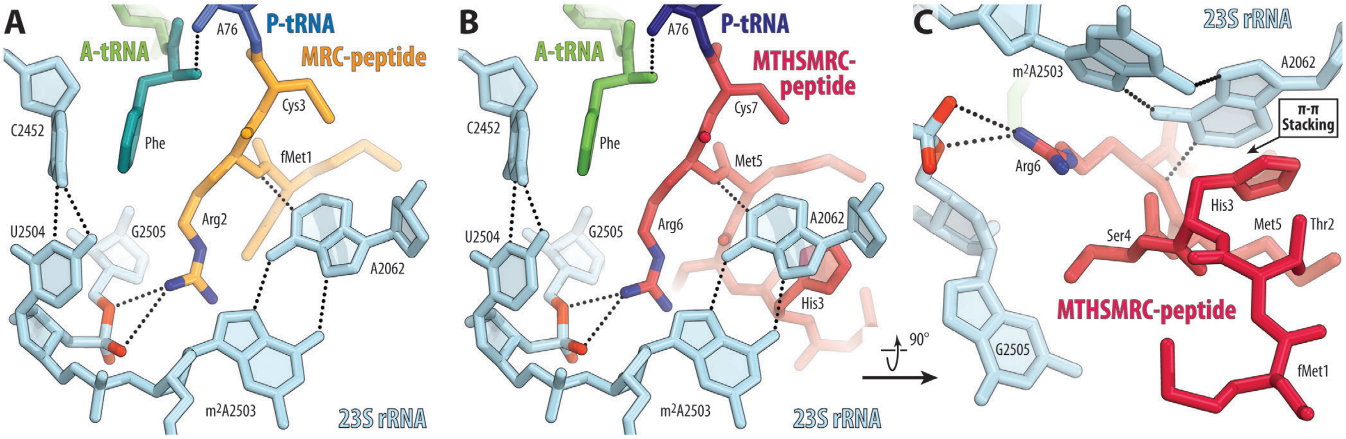


**Supplementary Figure 5. Additional interactions of the side chains of fMRC- and fMTHSMRC-peptidyl-tRNAs with the ribosome.** (**A, B**) Close-up views of the electrostatic interactions between the side chain of the penultimate Arg residue of the P-site fMRC-NH-tRNA$_i^{Met}$ (A, blue with peptide highlighted in orange) or fMTHMRC-NH-tRNA$_i^{Met}$ (B, navy with peptide highlighted in crimson) and the phosphate of nucleotide G2505 of the 23S rRNA. H-bonds are shown by black dotted lines. (**C**) Stacking interactions between the aromatic side chain of His3 of fMTHMRC-peptidyl-tRNA and A2062 nucleobase of the 23S rRNA.

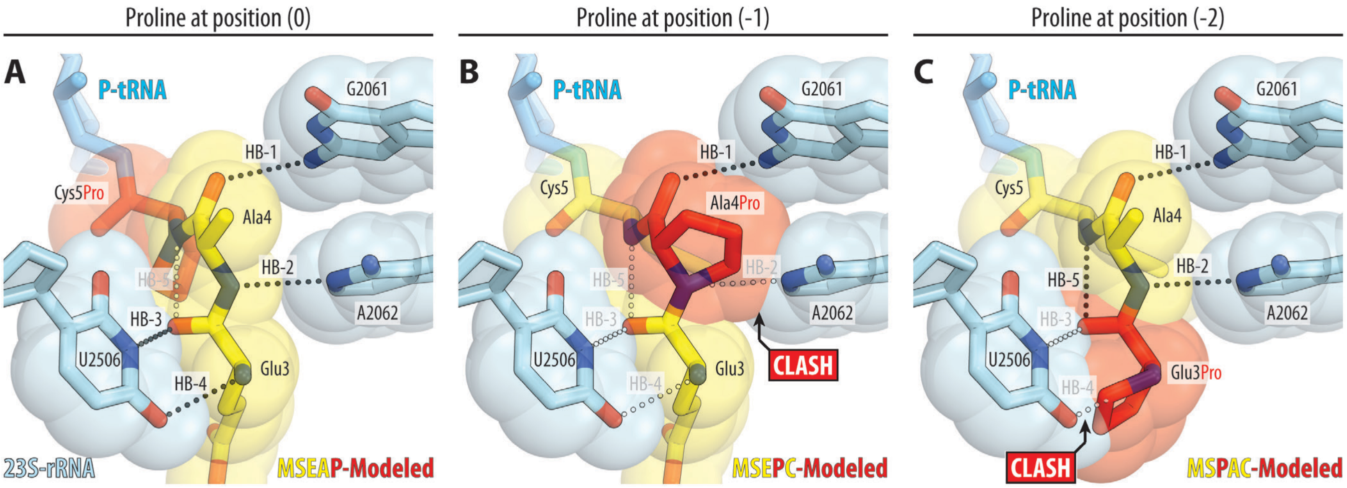


**Supplementary Figure 6. Proline residues in the nascent peptide are unable to form stabilizing H-bonds in the NPET.** (**A-C**) *In silico* modeling of proline residues at ultimate (A, Cys5Pro), penultimate (B, Ala4Pro), or pen-penultimate (C, Glu3Pro) positions of the fMSEAC peptide chain. Note that besides its inability to form most of the peptide-stabilizing H-bonds, proline in the penultimate and pen-penultimate positions clashes with nucleotides A2062 and U2506 of the 23S rRNA, respectively. Geometrically possible and impossible H-bonds are shown by black and white dotted lines, respectively.

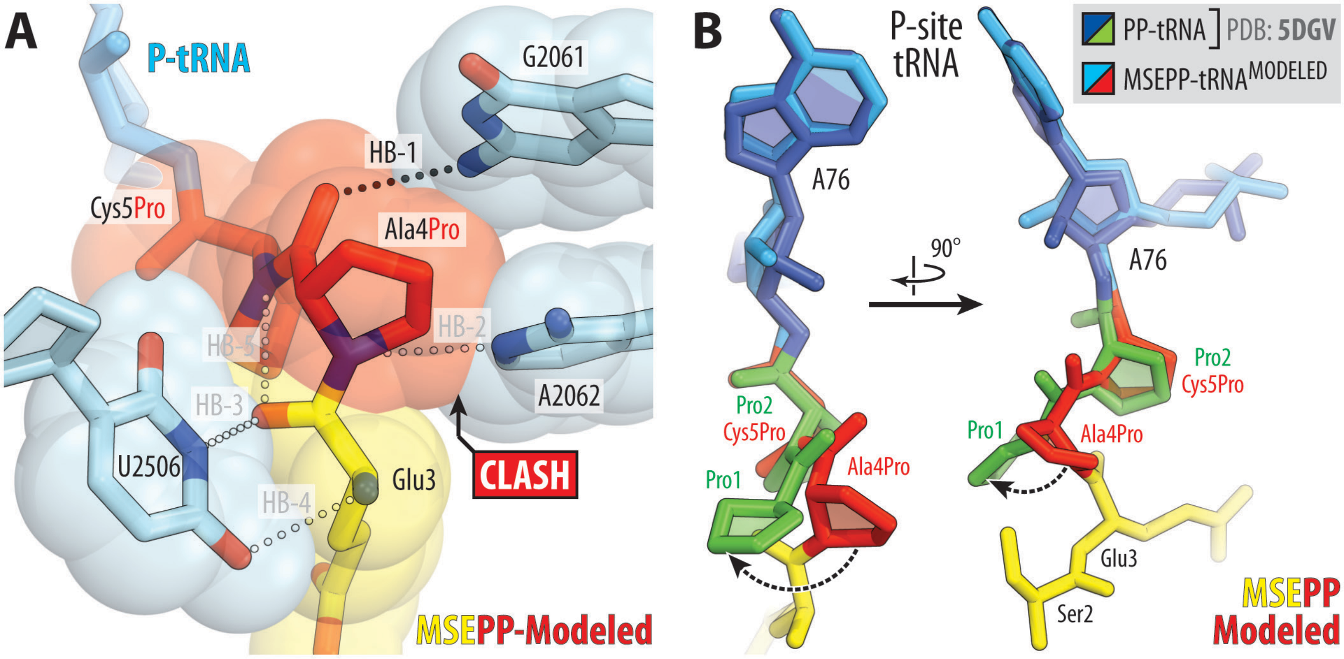


**Supplementary Figure 7. Proline residues alter the path of the nascent peptide in the NPET.** (**A**) *In silico* modeling of the two consecutive proline residues at ultimate (Cys5Pro) and penultimate (Ala4Pro) positions of the fMSEAC peptide chain. Note that, due to the side chains, the diproline-containing peptide cannot adopt a conformation possible for other peptides in the NPET and must re-orient. (**B**) Comparison of the previous structure of ribosome-bound short diprolyl-tRNA analog (green, PDB entry 5DGV (*12*)) with the *in silico*-modeled diprolyl-containing tRNA based on the structure of MSEAC-peptidyl-tRNA (red). Note that in order to avoid a steric clash with the A2062, the diprolyl moiety of the nascent peptide deviates to the side (black dashed arrows) and, thus, has an alternative path in the NPET.

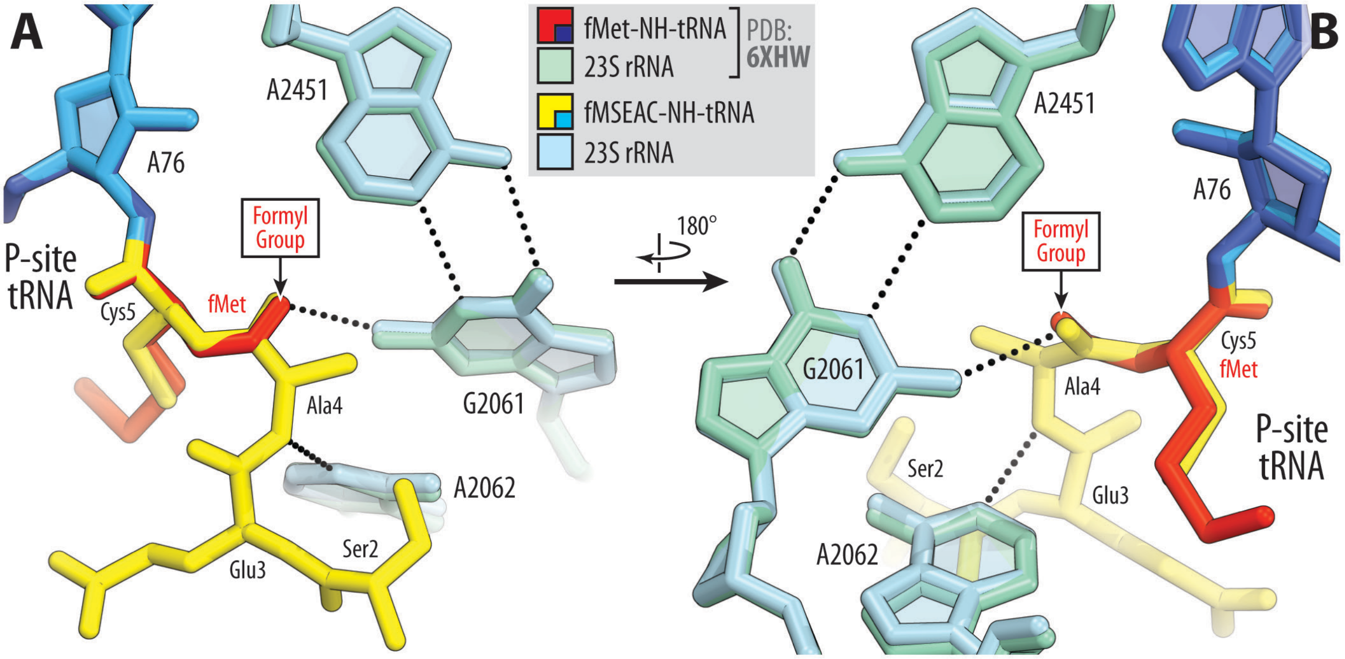


**Supplementary Figure 8. Formylation of the first methionine residue provides additional stability to the initiator tRNA substrate in the P site.** Superpositioning of the previous structure of ribosome-bound initiator fMet-NH-tRNA$_i^{Met}$ (navy with the fMet moiety in red, PDB entry 6XHW (*6*)) with the new structure of fMSEAC-peptidyl-tRNA (blue with the peptide moiety highlighted in yellow) viewed from two opposite sides (**A, B**). H-bonds are shown by black dotted lines. Note that the positions of carbon and oxygen atoms in the formyl group and those in the carbonyl group of the penultimate residue in the nascent peptide chain are nearly identical, ensuring formation of the same H-bond with the exocyclic amino group of the G2061 residue.
